# Pronounced α-synuclein pathology in a seeding-based mouse model is not sufficient to induce mitochondrial respiration deficits in the striatum and amygdala

**DOI:** 10.1101/2020.03.26.006189

**Authors:** Johannes Burtscher, Jean-Christophe Copin, Carmen Sandi, Hilal A. Lashuel

**Author notes:** To whom correspondences should be addressed, EPFL Station 19, CH-1015 Lausanne, Switzerland.

## Abstract

Increasing evidence suggests that crosstalk between α-synuclein pathology formation and mitochondrial dysfunctions plays a central role in the pathogenesis of Parkinson’s disease and related synucleinopathies. While mitochondrial dysfunction is a well-studied phenomenon in the substantia nigra, which is selectively vulnerable in Parkinson’s disease and some models thereof, less information is available in other brain regions that are also affected by synuclein pathology.

Therefore, we sought to test the hypothesis that early α-synuclein pathology causes mitochondrial dysfunction, and that this effect might be exacerbated in conditions of increased vulnerability of affected brain regions, such as the amygdala.

We combined a model of intracerebral α-synuclein pathology seeding with chronic glucocorticoid treatment modelling non-motor symptoms of Parkinson’s disease and affecting amygdala physiology. We measured mitochondrial respiration, ROS generation and protein abundance as well as α-synuclein pathology in male mice.

Chronic corticosterone administration induced mitochondrial hyperactivity in the amygdala. Although injection of α-synuclein preformed fibrils into the striatum resulted in pronounced α-synuclein pathology in both striatum and amygdala, mitochondrial respiration in these brain regions was altered in neither chronic corticosterone nor control conditions.

Our results suggest that early stage α-synuclein pathology does not influence mitochondrial respiration in the striatum and amygdala, even in corticosterone-induced respirational hyperactivity. We discuss our findings in light of relevant literature, warn of a potential publication bias and encourage scientist to report their negative results in the frame of this model.

**Significance statement:** We provide evidence that early stage synucleinopathy by itself or in combination with exogenous corticosterone induced amygdala hyperactivity did not compromise mitochondrial respiration in the striatum and amygdala in one of the most commonly used models of synucleinopathies. These results may explain, why this model in the hands of many research groups does not elicit pronounced Parkinson’s disease like symptoms in the absence of mitochondrial dysfunction in brain regions strongly affected by synuclein pathology and involved in non-motor (amygdala) and motor (striatum) symptoms. Our findings call for rigorous investigation of the short- and long-term effects of α-synuclein pathology on mitochondrial function/dysfunction in this model, in particular in brain regions strongly affected by neurodegeneration such as the substantia nigra pars compacta.

## Introduction

The misfolding, aggregation and accumulation of α-synuclein in Lewy bodies and selective neurodegeneration of dopaminergic neurons are defining hallmarks of Parkinson’s disease (PD), the most common neurodegenerative motor-disease (Spillantini et al., 1997; Lashuel et al., 2013). Mutations or multiplications of the α-synuclein encoding gene, *SNCA*, cause familial forms of PD (Polymeropoulos et al., 1997; Kruger et al., 1998; Singleton et al., 2003; Ibanez et al., 2004; Zarranz et al., 2004), thereby suggesting a causal role of α-synuclein in PD. Increasing evidence supports the Braak hypothesis of α-synuclein pathology propagation during the progression of PD (Braak et al., 2003b; Braak et al., 2003a); 1) α-synuclein pathology spreading from host PD patient brains to mesencephalic transplants grafted into these brains (Kordower et al., 2008; Li et al., 2008); 2) inter-cellular transmission of α-synuclein pathology (Desplats et al., 2009; Volpicelli-Daley et al., 2011); 3) α-synuclein pathology seeding in mouse brain using recombinant α-synuclein preformed fibrils (PFFs) or PD or multiple system atrophy brain-derived aggregates resulting in wide-spread brain pathology (Luk et al., 2012; Masuda-Suzukake et al., 2013; Recasens et al., 2014).

Mitochondrial dysfunction, in particular electron transport chain deficiencies, has also been implicated in PD pathogenesis (Langston et al., 1983; Schapira et al., 1989) and is thought to play important roles in both neurodegeneration and α-synuclein pathology (Hsu et al., 2000). Aggregated forms of α-synuclein has been shown to interfere with the regulation of mitochondrial import (Di Maio et al., 2016), mitochondrial membrane potential, reactive oxygen species (ROS)-generation and morphology (Tapias et al., 2017; Grassi et al., 2018) and oxidative phosphorylation (Ludtmann et al., 2016; Ludtmann et al., 2018) (Devi et al., 2008; Mahul-Mellier et al., 2020). However, whether α-synuclein pathology in the brain is sufficient to precipitate mitochondrial dysfunction is not known. We hypothesized that models of α-synuclein pathology seeding provide unique opportunities to address this question. α-synuclein pathology seeding in rodents by inoculation of the striatum (Luk et al., 2012) or the olfactory bulb (Rey et al., 2016) with PFFs results in strong α-synuclein pathology in the amygdala (Burtscher et al., 2019), reminiscent of human synucleinopathy (Nelson et al., 2018; Sorrentino et al., 2019). Recently, we reported subtle behavioral effects related to this pronounced α-synuclein pathology in the mouse amygdala (Burtscher et al., 2019), with little influence of additional exogenous corticosterone induced amygdala-related mood deficits, modelling prodromal mood symptoms in PD. Metabolic alterations in the amygdala have been reported upon chronic glucocorticoid administration (Thobois et al., 2017), thus we hypothesized that mitochondrial dysfunction might precede behavioral deficits. Therefore, we assessed, whether α-synuclein pathology at early time points (5-6 weeks) after seeding in the amygdala would suffice to induce mitochondrial dysfunction and whether additional chronic corticosterone would further exacerbate mitochondrial dysfunction. We chose this time point because this is when we see maximum α-synuclein pathology (pS129 immunoreactivity) in the amygdala (Burtscher et al., 2019). Originally, continuously increasing α-synuclein pathology in the amygdala has been assumed after injection in the striatum (Luk et al., 2012) or olfactory bulb (Rey et al., 2016).

The phenomenon of dropping pS129 immunoreactivity in limbic and cortical circuits, although potentially strain dependent, is increasingly acknowledged by some (Rey et al., 2019) but not other groups (Henderson et al., 2019).

We hypothesized that the peaking α-synuclein pathology in the amygdala 1 to 3 months after PFF injection would induce mitochondrial dysfunction contributing to disease progression. We did not investigate time points and brain regions with clear neuronal loss to exclude potential confounding with cellular loss induced mitochondrial dysfunction.

## Methods

### PFFs

α-synuclein fibrils were prepared and characterized as described previously (Burtscher et al., 2019; Kumar et al., 2020). Brieflly, a 325 μM solution of pure mouse α-synuclein recombinant protein (in PBS, pH7.2) was centrifuged in filter tubes of 0.2 μM (5 min, 5000 rpm). Supernatant was shaken at 900 rpm (4 days, 37°C). Resulting PFFs were sonicated (one 1 s pulse per 2 s for 10 s at 40% amplitude) and stored at −80 °C until use.

For characterization samples were treated with 10 μM Thioflavin T (ThT, in 50 mM glycine, pH8.5) in black 384-well plates (Nunc) and the extent of fibril formation was measured at 485 nm, excitation 450 nm using a Bucher Analyst AD plate reader.

To assess the amount of fibrils formed and monomer release we used sedimentation (100000g, 30 min) and filtration assays (14000 g for 15 min through a 100 kD filter). Electron microscopy was applied for morphological characterizations.

### Animals

C57BL/6JRj male mice (2-3 months of age, 3 animals per cage) were housed at 23 °C, 40 % humidity, light from 7am-7pm and dark from 7pm-7am with free access to standard laboratory rodent chow and water for in-vivo experiments. Primary hippocampal cultures were derived from P0-2 pups of C57BL/6JRccHsd mice.

All animal experimentation procedures were approved by the Cantonal Veterinary Authorities (Vaud, Switzerland) and performed in compliance with the European Communities Council Directive of 24 November 1986 (86/609EEC). Every effort was taken to minimize number and stress of the animals.

5 μg of PFFs in 2 μL PBS were injected into the right dorsal striatum (AP +0.4, ML +2, DV - 2,6) of fully anesthetized animals (100 mg/kg ketamine and 10 mg/kg xylazine, i.p.) on stereotactic frames (Kopf Instruments) through a 34-gauge canula using a 10 μL Hamilton syringe (flow rate of 0.1 ul/min).

Mice were delivered an overdose of pentobarbital (150 mg/kg) for transcardial perfusion with heparinized saline followed by 4 % paraformaldehyde for histology (N=3 per group, total N=24). Mice for mitochondrial respiration studies (N=9 per group, N=36) were sacrificed by neck dislocation and exsanguination and tissues were directly dissected and prepared for high resolution respirometry.

### Primary neuronal cultures, PFF treatment and immunocytochemistry

Primary hippocampal neurons were prepared from P0-2 pups of C57BL/6JRccHsd mice (Harlan) and imaged as described before (Mahul-Mellier et al., 2020). Briefly, hippocampi were dissescted in Hank’s Balanced Salt Solution (HBSS) and digested by papain (20U/mL, Sigma-Aldrich) for 30 min at 37 ◻C. Papain activity was inhibited using a trypsin inhibitor (Sigma-Aldrich) and tissues were dissociated by trituration. The cells were resuspended in adhesion media (MEM, 10% Horse Serum, 30% glucose, L-glutamine and penicillin/streptomycin) (Life Technologies) and plated in 6-well plates coated with poly-L-lysine 0.1% w/v in water (Brunschwig) at a density of 250,000 cells/ml. Adhesion media was replaced by Neurobasal medium (Life Technologies) containing B27 supplement (Life Technologies), L-glutamine and penicillin/streptomycin (100U/mL, Life Technologies) after 3 h. After 5 days in vitro, primary cultures were treated with 70 nM of PFFs. 14 days later cells washed in PBS, fixed in 4% PFA for 20 min and immunostained For mitotracker experiments, cells were exposed to 100 nM MitoTracker Red CMXRos (Invitrogen) 30 min before fixation. For immunostaining, fixed cells were blocked in 3 %BSA in 0.1% Triton X-100 PBS for 30 min, exposed to primary antibodies for 2 h, washed and incubated with secondary antibodies (plus DAPI) for 1 h, washed and mounted using Fluoromount.

### Assessment of body composition

Fat- and lean-mass was measured by Echo MRI before (to ensure similar body composition across experimental groups) and 8 weeks after corticosterone/vehicle treatment.

### Continuous exogenous corticosterone treatment

Corticosterone (Sigma) was dissolved in 0.45 % hydroxypropyl-b-cyclodextrin (Sigma) and administered (35 mg/l) to animals in drinking water continuously starting 4 weeks before surgery until sacrifice of the animals as described previously (Bacq et al, 2012). Control animals were administered vehicle (0.45 % hydroxypropyl-b-cyclodextrin) in the same manner.

### Histology and immunohistochemistry

Perfused brains were postfixed in 4% paraformaldehyde, embedded in paraffin and cut coronally to 4 μm. Brain slices were de-waxed and epitope retrieval for 20 min at 95°C in trisodium citrate buffer (10 mM, pH 6.0) in a retriever (Labvision) was applied. For immunofluorescence, they were blocked for 60 min in 3% bovine serum albumin in PBS containing 0.1% Triton X-100. Primary antibodies were applied over night at 4°C and secondary antibodies (plus DAPI) for immunofluorescence for 60 min at RT, before mounting the slides using fluoromount. Sections of N=3 animals per time point (30 dpi and 360dpi of only PFFs or PBS, total N=12) were used for pS129 stainings together with ubiquitin, p62 or Thioflavin S (ThS). For ThS stainings, sections were incubated for 15 min in 0.01% ThS and washed in 80% ethanol, followed by washing in water and then PBS before blocking and immunostaining.

Sections of N=3 animals per group (total N=12) at 60 dpi (corticosterone / vehicle and PFF / PBS) were used for stainings for pS129, N-terminal (1-20) α-synuclein and TOM20, with or without proteinase K treatment (8 min in 1 μg/mL of proteinase K in 50mM TrisHCl buffer at pH 7.4 before staining). Imaging was performed with a Zeiss LSM700 confocal microscope.

Other sections of the same animals were used for DAB-stainings and exposed to 3% H_2_O_2_ in PBS for 30 min before blocking for 60 min in 3% bovine serum albumin in PBS containing 0.1% Triton X-100 at RT. Primary antibody against α-synuclein pS129 (Wako Chemicals USA, 014-20281, 1:10000) was applied over night at 4°C, and ImmPRESS reagent anti-mouse IgG was applied for 40 min at RT followed by incubation for 10 min in 3,3’-diaminobenzidine (DAB) dissolved in 50 mM Tris buffer and 0.06% H_2_O_2_. Sections were counterstained with Mayer’s hematoxylin, mounted with fluoromount an imaged using an Olympus AX70 microscope.

### Respirometry

After sacrifice, striatum and amygdala-enriched tissues (both from hemisphere of injection and contralateral hemisphere) were dissected on ice using a mouse brain matrix (Agnthos). Wet tissue was weighed and collected in ice-cold BIOPS (2.8 mM Ca_2_K_2_EGTA, 7.2 mM K_2_EGTA, 5.8 mM ATP, 6.6 mM MgCl_2_, 20 mM taurine, 15 mM sodium phosphocreatine, 20 mM imidazole, 0.5 mM dithiothreitol and 50 mM MES, pH = 7.1), homogenized in ice-cold MiR05 (0.5 mM EGTA, 3mM MgCl_2_, 60 mM potassium lactobionate, 20 mM taurine, 10 mM KH_2_PO_4_, 20 mM HEPES, 110 mM sucrose and 0.1% (w/v) BSA, pH = 7.1) using a pestle for eppendorf tubes in a concentration of 1 mg wet-weight per 10 μL MiR05. Respiration was measured in parallel to mitochondrial ROS production (O_2_^−^ and H_2_O_2_) at 37 °C in the Oroboros O2k equipped with the O2K Fluo-LED2 Module (Oroboros Instruments, Austria). For mitochondrial ROS-measurement LEDs for green excitation were applied and a concentration of 1 mg wet tissue per ml MiR05 was added to final concentrations of 10 μM amplex red, 1 U/ml horse radish peroxidase and 5 U/ml superoxide dismutase in 2 ml MiR05 per O2K chamber. Calibration was performed by titrations of 5 μL of 40 μM H_2_O_2_.

Respirational states were assessed using a standard high resolution respirometry protocol: NADH-pathway respiration in the LEAK (N_*L*_) state was initiated using malate (2mM), pyruvate (10mM) and glutamate (20mM). Oxidative phosphorylation (N_*P*_) was stimulated by ADP (5 mM). Succinate (10 mM) addition yielded NADH- and Succinate-linked respiration in oxidative phosphorylation (NS_P_) and in the uncoupled state (NS_*E*_) after titration (∆0.5 μM) of carbonyl cyanide m-chlorophenyl hydrazine (CCCP). Complex I inhibition by rotenone (0.5 μM) allowed analysis of succinate-linked respiration in the uncoupled state (S_*E*_). All oxygen fluxes were corrected for residual (non-oxidative phosphorylation associated) oxygen consumption, ROX, after antimycin A addition. Respiratory acceptor control ratios (RCR) were calculated as N_*L*_/NS_*P*_. Flux control ratios (FCRs) were assessed of NADH-(N_*P*_ / NS_*E*_) succinate-driven respiration (S_*E*_ / NS_*E*_). Mitochondrial ROX-values were corrected for background and are presented as mitochondrially generated O_2_ ^−^ and H O per mg wet weight and second, for the respective states.

### Western blotting and fractionated sample preparations

After using the required volume for respirometry, phosphatase inhibitor mixes (Sigma, P5726 and P0044) and protease inhibitor mix (Sigma, P8340) at 1:100 and 1 mM phenylmethylsulfonyl fluoride (PMSF) were added to tissue homogenates, the samples were snap-frozen in liquid nitrogen and stored at −80°C. For extraction of Triton X-100 soluble and insoluble fractions, homogenates were diluted 1:1 in 1% Triton X-100/ Tris-buffered saline (TBS) (50 mM Tris, 150 mM NaCl, pH 7.5) including protease and phosphatases inhibitor mixes and PMSF at the same concentration as indicated above (total fractions). They were sonicated 10x at a 0.5s pulse (20% amplitude, Sonic Vibra Cell, Blanc Labo), incubated on ice for 30 min and centrifuged at 100,000 g (30 min, 4°C). The supernatant was used as soluble fraction and the pellet was washed in 1% Triton X-100/TBS, sonicated as above, and centrifuged again at 100,000 g (30 min, 4°C). The pellet (insoluble fraction) was resuspended in 2% sodium dodecyl sulfate (SDS)/TBS including protease and phosphatases inhibitor mixes and PMSF at the same concentration as indicated above, and sonicated 15x at a 0.5s pulse (20% amplitude). While sufficient tissue for tissue extractions were available for striatal samples (N=8-9 per condition), 2-3 amygdala samples were pooled (at least N=3 per condition).

Protein concentrations of different fractions were assessed using a BCA assay. 15-20 μg of proteins were loaded on a 16% tricine gel, transferred onto a nitrocellulose membrane (Fisher Scientific, Switzerland) using a semi-dry system (Bio-Rad, Switzerland). 30◻min of blocking in Odyssey blocking buffer (Li-Cor Biosciences, Bad Homburg, Germany) was followed by incubation probed overnight at 4◻°C with primary antibodies. Membranes were then washed in 0.01% (v/v) Tween-20 (Sigma-Aldrich) in PBS (PBS-T), and secondary antibodies were applied for 1h at RT. After final washing in PBS-T, membranes were scanned using a Li-COR scanner (Li-Cor Biosciences).

### Antibodies

Antibodies applied in this study are listed in table 1.

**Table 1.**
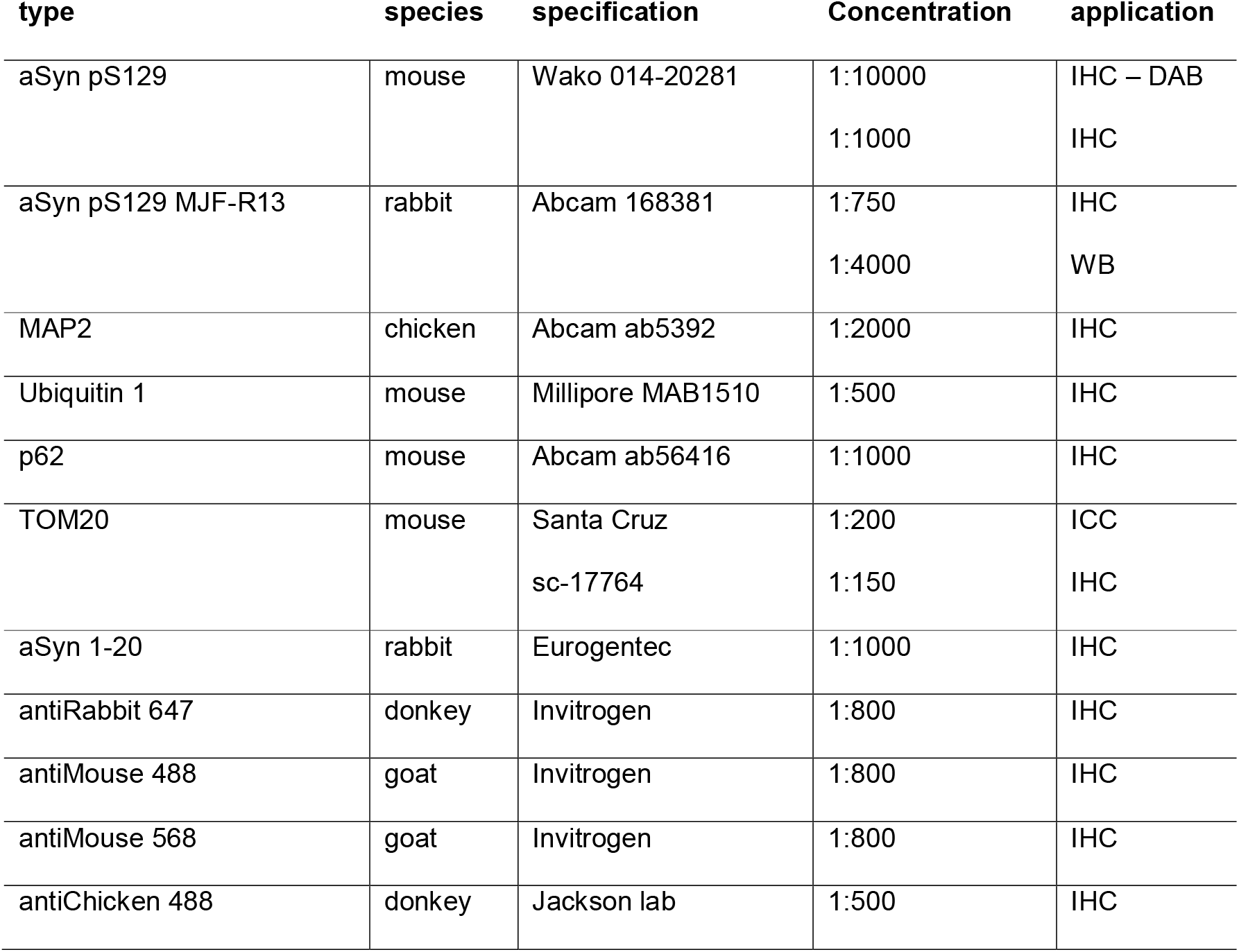
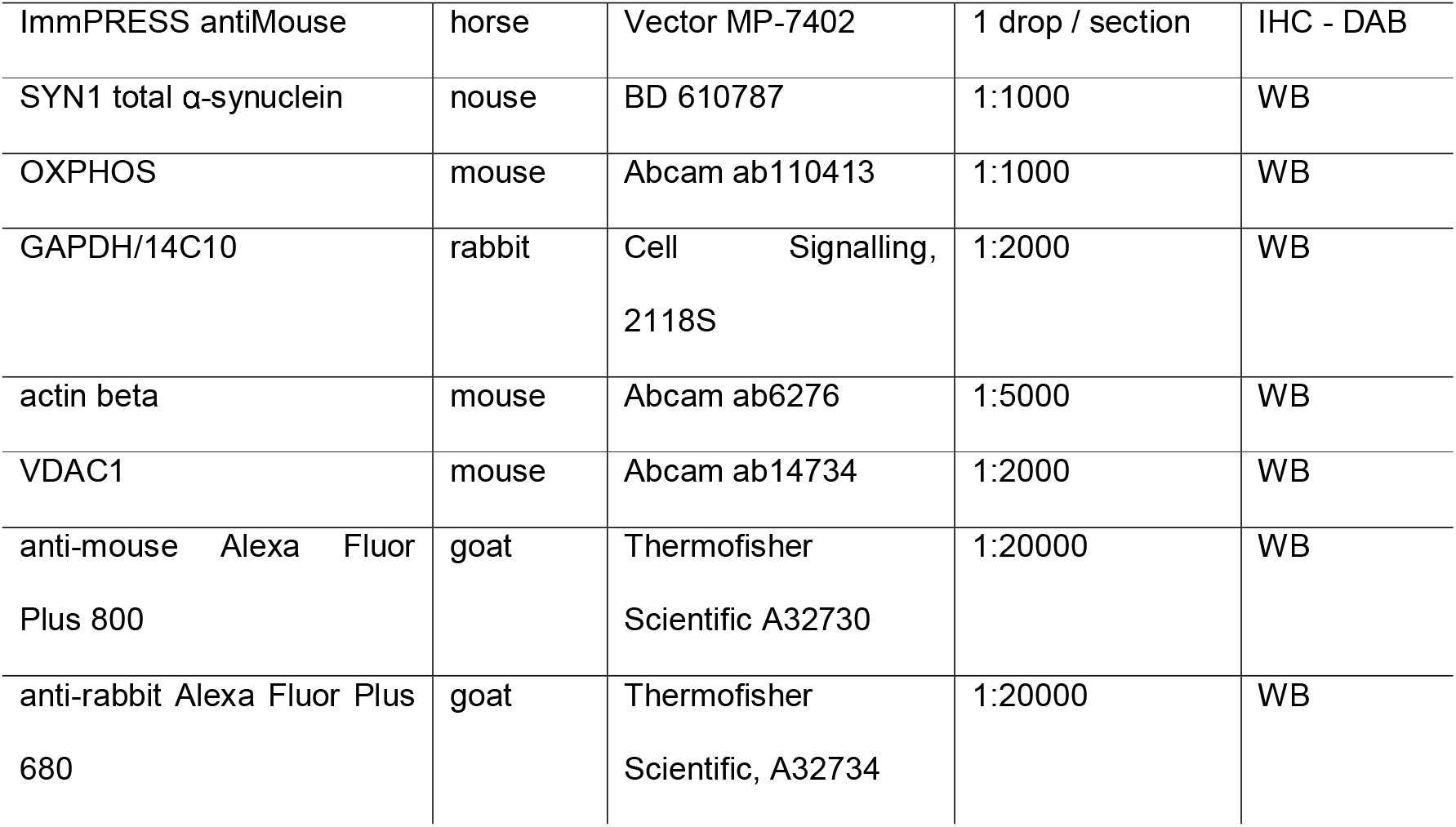
antibodies. IHC – immunohistochemistry, ICC – immunocytochemistry, WB – western blot

| type | species | specification | Concentration | application |
| --- | --- | --- | --- | --- |
| aSyn pS129 | mouse | Wako 014-20281 | 1:10000 | IHC – DAB |
|  |  |  | 1:1000 | IHC |
| aSyn pS129 MJF-R13 | rabbit | Abcam 168381 | 1:750 | IHC |
|  |  |  | 1:4000 | WB |
| MAP2 | chicken | Abcam ab5392 | 1:2000 | IHC |
| Ubiquitin 1 | mouse | Millipore MAB1510 | 1:500 | IHC |
| p62 | mouse | Abcam ab56416 | 1:1000 | IHC |
| TOM20 | mouse | Santa Cruz | 1:200 | ICC |
|  |  | sc-17764 | 1:150 | IHC |
| aSyn 1-20 | rabbit | Eurogentec | 1:1000 | IHC |
| antiRabbit 647 | donkey | Invitrogen | 1:800 | IHC |
| antiMouse 488 | goat | Invitrogen | 1:800 | IHC |
| antiMouse 568 | goat | Invitrogen | 1:800 | IHC |
| antiChicken 488 | donkey | Jackson lab | 1:500 | IHC |
| ImmPRESS antiMouse | horse | Vector MP-7402 | 1 drop / section | IHC - DAB |
| SYN1 total α-synuclein | nouse | BD 610787 | 1:1000 | WB |
| OXPHOS | mouse | Abcam ab110413 | 1:1000 | WB |
| GAPDH/14C10 | rabbit | Cell Signalling,<br>2118S | 1:2000 | WB |
| actin beta | mouse | Abcam ab6276 | 1:5000 | WB |
| VDAC1 | mouse | Abcam ab14734 | 1:2000 | WB |
| anti-mouse Alexa Fluor<br>Plus 800 | goat | Thermofisher<br>Scientific A32730 | 1:20000 | WB |
| anti-rabbit Alexa Fluor Plus<br>680 | goat | Thermofisher<br>Scientific, A32734 | 1:20000 | WB |

### Statistics

Corticosterone versus vehicle and PFF injections versus PBS injections were compared in a 2×2 statistical design. 2way ANOVAs were calculated, if Gaussian normality could be assumed. Normality was tested using the Anderson-Darling, D’Agostino-Pearson omnibus, Shapiro-Wilk and Kolmogorov-Smirnov tests.

All absolute values and western blot values are presented as mean + SD. Normalized respirational and mitochondrial ROS values are presented as mean + SEM.

## Results

### Mitochondrial coupling efficiency is maintained in the presence of both corticosterone induced alteration of body composition and seeded α-synuclein pathology

First, we verified the effects of chronic corticosterone and injection of α-synuclein preformed fibrils (PFFs) (Fig. 1A). Similar to our recent report (Burtscher et al., 2019), chronic corticosterone increased body fat tissue content, as assessed by Echo MRI (F_*corticosterone*_(1,31) = 69,45, p<0,0001, Fig. 1B) independently of PFF-injection. α-Synuclein pS129 pathology was confirmed by histology in ipsilateral striatum and amygdala (Fig. 1C). We observed similar pathology patterns in PFF-groups, irrespective of corticosterone treatment as reported previously (Burtscher et al., 2019).

**Fig. 1:**
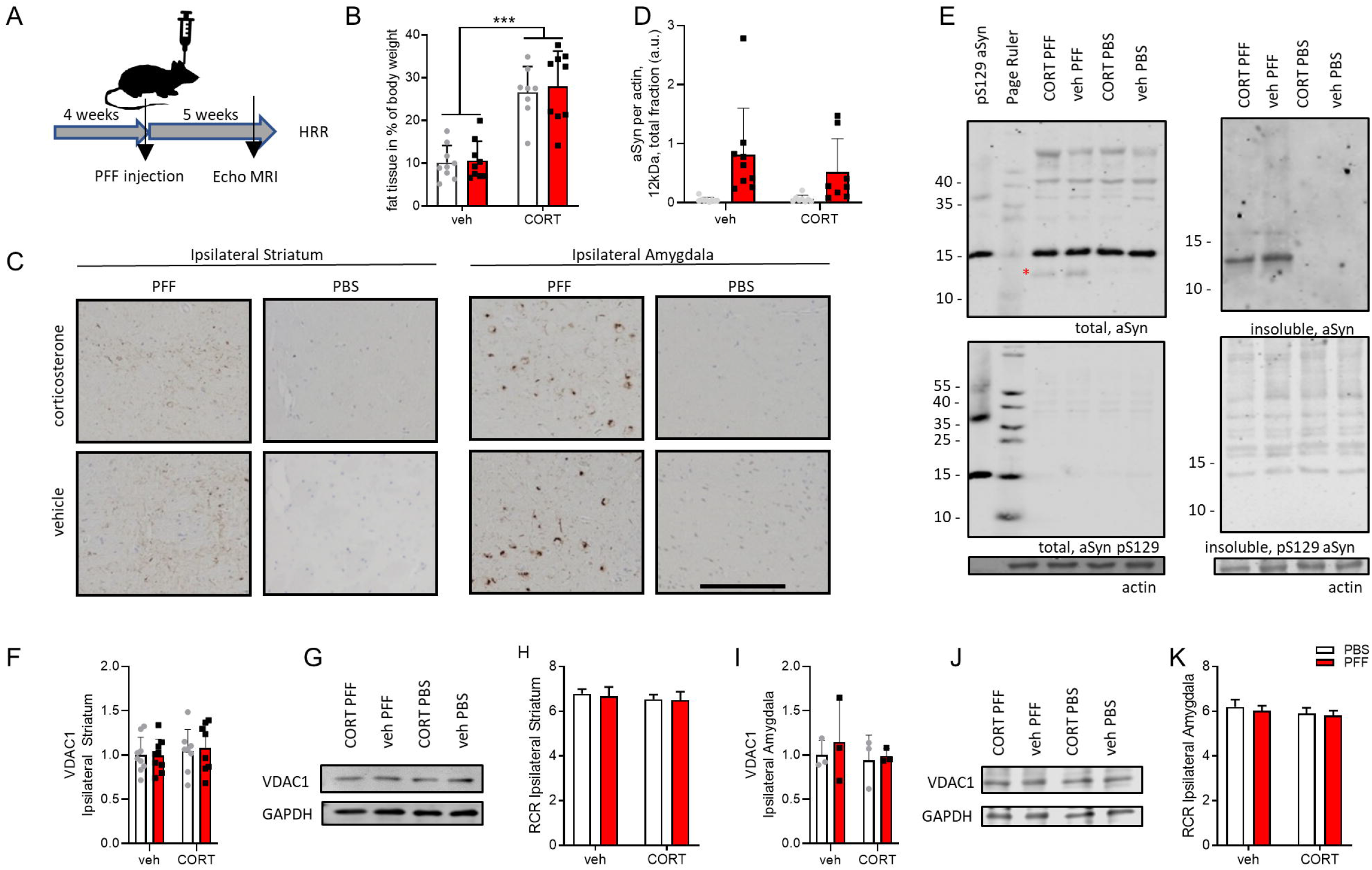
Effects of chronic corticosterone treatment and of α-synuclein (α-synuclein) inocculation. Animals were treated with corticosterone (CORT) or vehicle (veh) in the drinking water for 1 month, after which they were injected unilaterally into the dorsal striatum with 5 μg PFFs or solvent (PBS). CORT/veh treatment was continued until around 5 weeks after surgery, when the animals were sacrificed (A). Animals had massively increased fat tissue as assessed by echo MRI after 8 weeks of CORT treatment (B). 6 weeks after PFF injection clear α-synuclein pathology (as evidenced here by α-synuclein pS129 immunostaining) was observed both in the striatum (mainly neuritic) and in the amygdala (mainly somatic)(C). Striatum and amygdala were dissected for high resolution respirometry and residual tissues were used for biochemical analyses. PFF-injected striatum contained truncated α-synuclein (running at 12kDa) in total homogenate, as quantified in (D) and represented in (E, asterisk indicates truncated 12 kDa α-synuclein), but even more clearly in insoluble fractions (D). VDAC1 protein levels and mitochondrial respiratory control ratios (RCRs) were similar across groups both in striatum (F, G, H) and amygdala (I, J, K). *** indicates p<0.001. pS129 α-synuclein label for western blot lanes denotes recombinant protein controls. Scale bar in (E) is 200 μm.

Previous studies have shown that α-synuclein is subjected to proteolytic processing during or after its aggregation that lead to both C- and N-terminal truncation of the protein (Baba et al., 1998; Gai et al., 1999; Mahul-Mellier et al., 2018{Grassi, 2018 #280)}. Therefore, western blot analysis was performed to investigate α-synuclein processing both in striatum and amygdala 5-6 weeks after striatal PFF-injection. We observed truncated forms (running at 12 kDa), a signature of pathological α-synuclein (Mahul-Mellier et al., 2019), in total striatal fractions of all PFF-conditions ipsilaterally (hemisphere of injection, Fig. 1D), although with high variation. Virtually all of the α-synuclein detected in all insoluble ipsilateral striatal fractions of PFF-but not PBS-conditions was truncated (Fig. 1E). Full length α-synuclein (15 kDa) levels were similar in total striatal fractions. In fractionated amygdala samples no or very little truncated α-synuclein was detected even in samples of PFF-injected mice. No α-synuclein pS129 signal was detected in any fraction by western blotting, maybe due to dilution of the locally restricted pathological tissue.

Next, we assessed general mitochondrial parameters, mitochondrial protein levels and coupling efficiency. VDAC1-levels, as well as mitochondrial coupling efficiencies, estimated by mitochondrial respiratory control ratios (RCRs) were similar across groups both in striatum (Fig. 1F, G, H) and amygdala (Fig, 1I, J, K).

Immunohistochemical analyses demonstrate increasing immunoreactivity of pS129 positive α-synuclein with Ubiquitin, Thioflavin S and p62 over time in the striatum (Fig. 2A) and amygdala (Fig. 2B). In primary hippocampal neurons we furthermore detected strong colocalization of pS129 with mitochondrial markers (Fig. 2C and (Mahul-Mellier et al., 2020)), which also was the case for neurons in the amygdala 60 dpi (Fig. 2D), which contained proteinase K resistant α-synuclein aggregations (Fig. 2E).

**Fig. 2:**
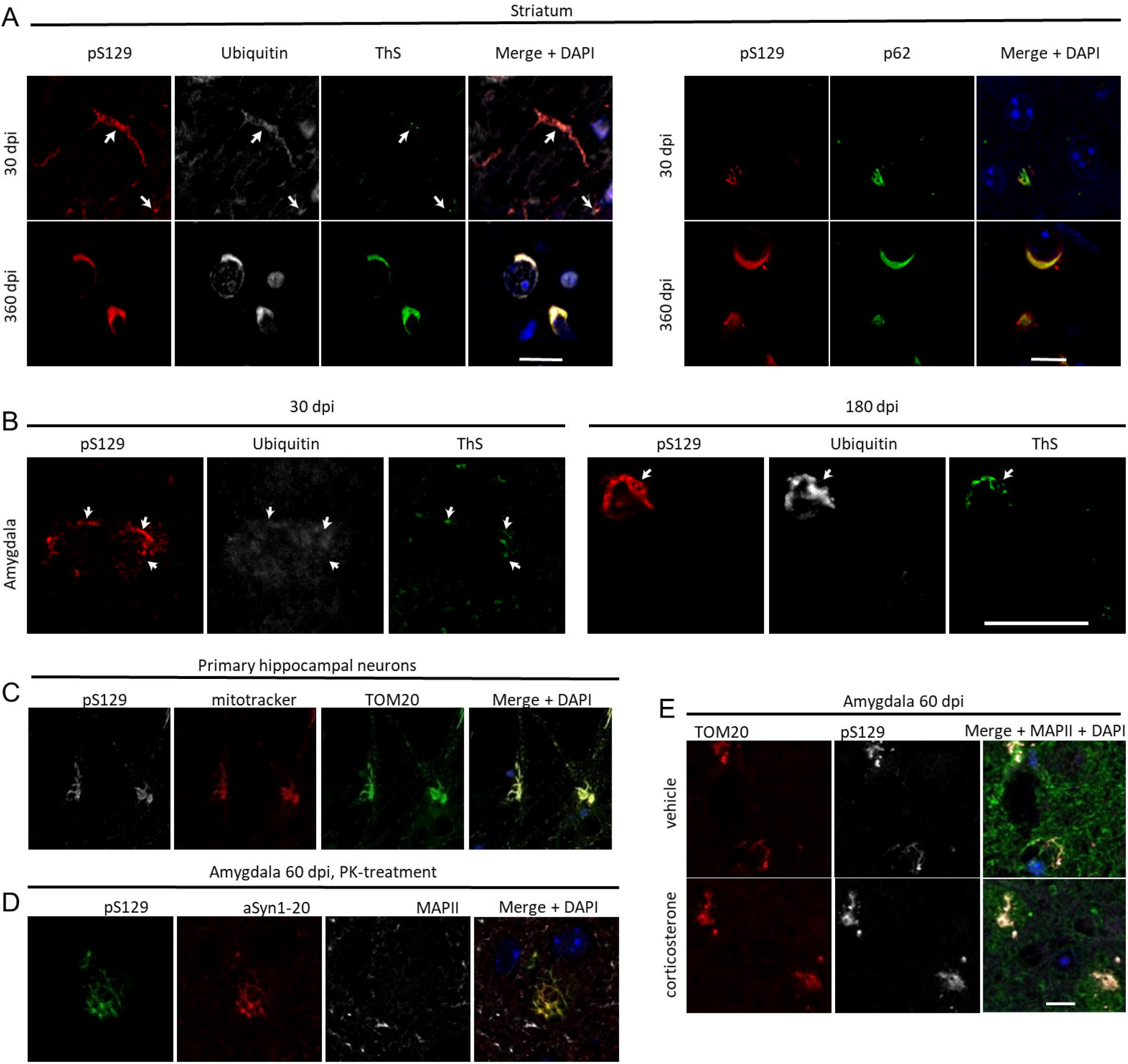
Characterization of α-synuclein (α-synuclein) pathology. In the striatum pS129 immunoreactivity at 30 days post injection (dpi) colocalized in peripheral cell compartments with ubiquitin, moderately with Thioflavin S (ThS) and p62. The colocalization was pronounced and perinuclear at 360 dpi (A). In the amygdala, similar colocalizations were observed at 30 and even more pronounced at 180 dpi (B). 14 days after treatment with 70 nM of PFFs, α-synuclein pathology colocalized strongly with the mitochondrial markers TOM20 and mitotracker in primary hippocampal neurons (C), with similar colocalizations being observed *ex-vivo* 60 dpi in the amygdala both in the chronic corticosterone and control condition (D). The aggregates were resistant to proteinase K at this time point in the amygdala (E). Green in the merged image in (D) corresponds to MAPII staining, and blue is DAPI staining. Scale bars in (A) are 10 μm, in (B) 20 μm and scale bar in (D) is also valid for (C) and (E) and is 10 μm.

### Corticosterone treatment, but not α-synuclein pathology alters mitochondrial respiration in amygdala

Having validated that the applied treatments induced the expected phenotypes, high resolution respirometry was applied concomitantly with amplex red fluometry to assess mitochondrial respiration and ROS production of striatum and amygdala tissues. In presence of nicotinamide adenine dinucleotide (NADH)-linked substrates and absence of adenosine diphosphate (ADP; no oxidative phosphorylation, LEAK-state N_*L*_), respiration in the amygdala (Fig. 3A), but not in the striatum (Fig. 3B) was increased in corticosterone treated mice. Similarly, respiration was higher in the corticosterone conditions in the NADH-driven oxidative phosphorylation (saturating concentrations of ADP, N_*P*_)(Fig. 3C,D), NADH- and succinate-driven oxidative phosphorylation (Fig. 3E,F, NS_*P*_) and NADH- and succinate-driven uncoupled (Fig. 3G,H, NS_*E*_) states in the amygdala (Fig. 3C,E,G), but not in the striatum (Fig. 3D,F,H). In the succinate-driven, uncoupled state after complex I inhibition (S_*E*_), differences of respiration were neither observed in the amygdala (Fig. 3I) nor striatum (Fig. 3J). Absolute respiration (per wet weight) and mitochondrial ROS production per brain region, as well as normalizations to VDAC1- or protein levels and statistical analyses are shown in table 2.

**Table 2.**
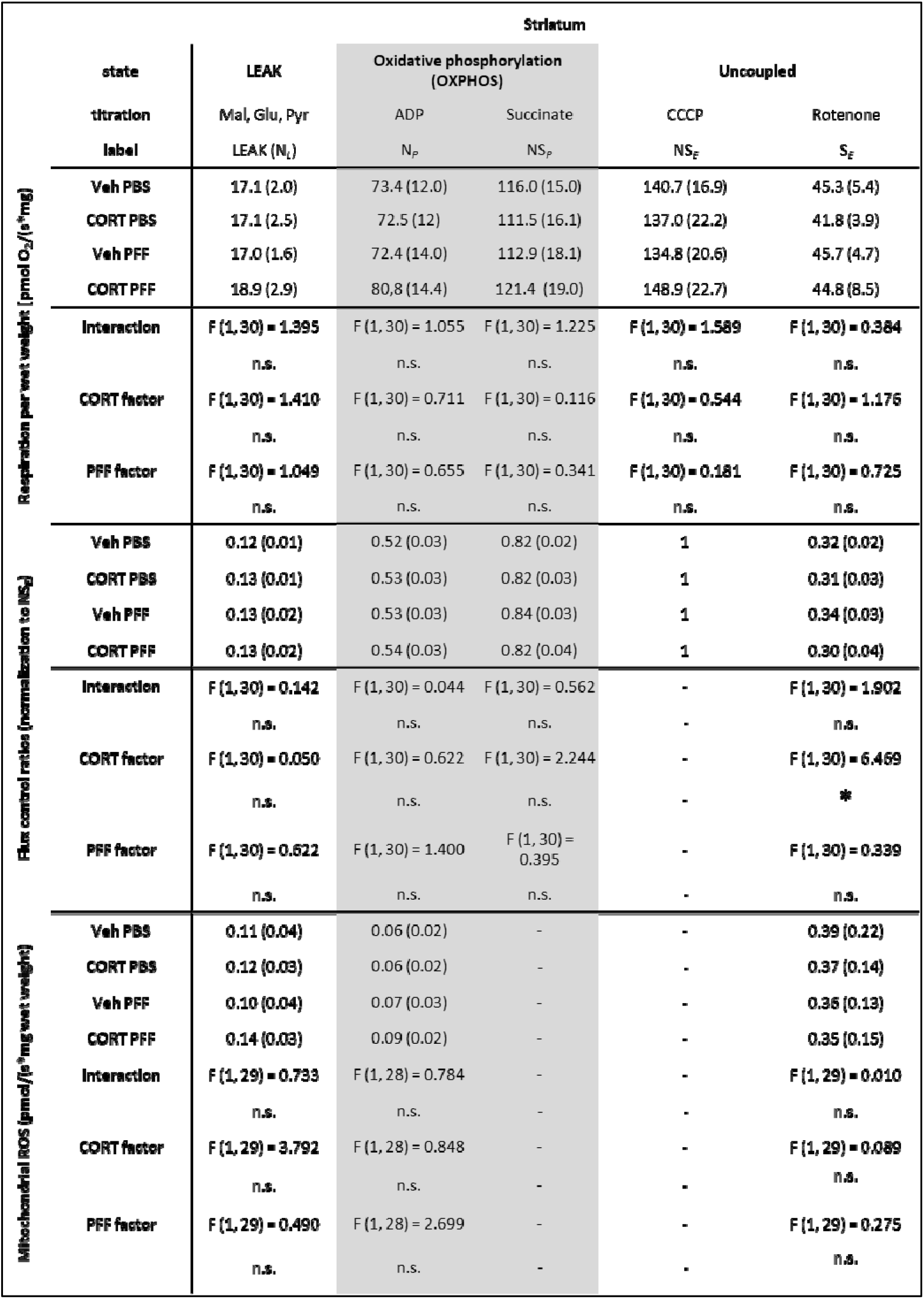
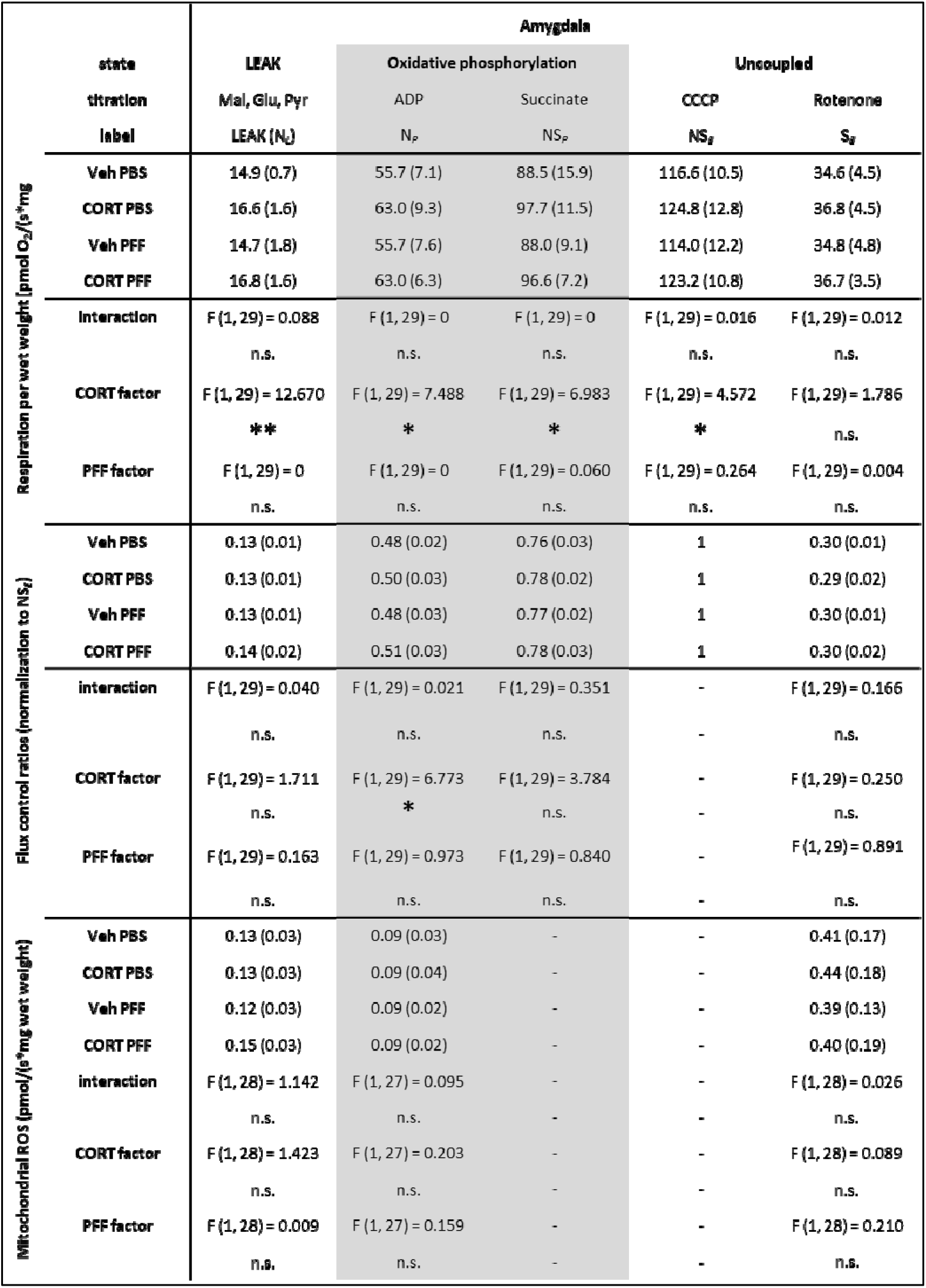
summary of high resolution respirometry data. * p<0.05, **p<0.01

**Fig. 3:**
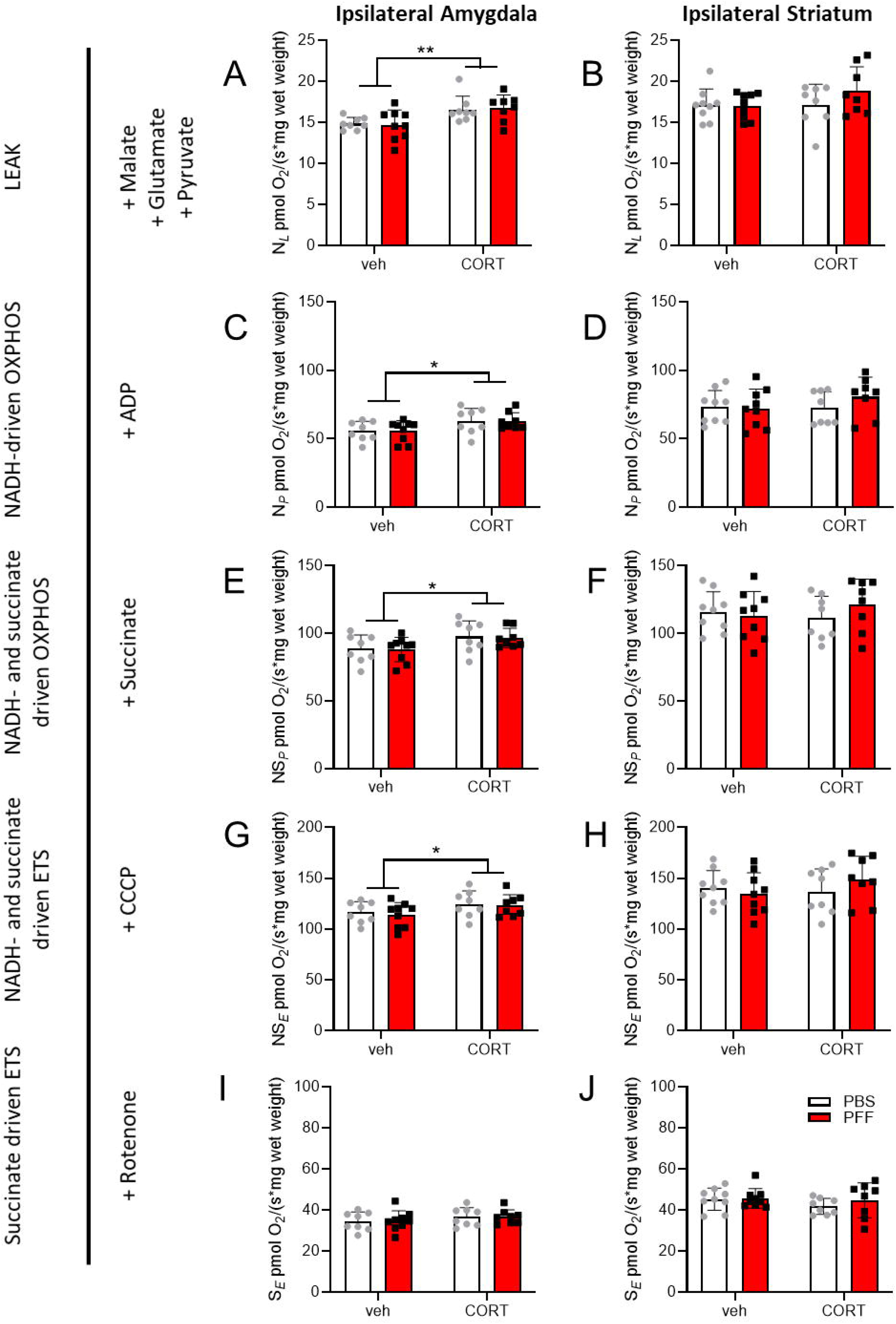
Testing mitochondrial functions *ex-vivo*. Striatum and amygdala were dissected for high resolution respirometry 9 weeks after corticosterone (CORT) or vehicle (veh) treatment, and/or after intrastriatal injection of 5 μg PFFs or solvent (PBS). Data from hemispheres of injections (=ipsilateral) are depicted. LEAK state in the presence of NADH substrates (N_L_) was measured (A, B), followed by addition of ADP yielding NADH-driven oxidative phosphorylation (N_P_) in (C, D), succinate to drive oxidative phosphorylation via both complex I and II (NS_P_) (E, F), CCCP to achieve the electron transport system maximum capacity (ETS, NS_E_) (G, H). Subsequent inhibition of complex I by rotenone resulted in succinate-driven, uncoupled respiration (S_E_) (I, J). Respirational values of all states are given for amygdala and striatum (hemisphere of injection = ipsilateral). Table to the right indicates respirational state and substance additions according to the SUIT (Substrate, Uncoupler, Inhibitor Titration) protocol. Main effects for CORT are indicated: * p<0.05, **p<0.01

To assess qualitative changes in respiration control, flux control ratios (FCRs) were calculated. While there were no differences in NADH-driven FCR (Fig. 4A), FCR was reduced for the uncoupled, succinate-driven state in corticosterone conditions (F_*corticosterone*_(1,30) = 6,634, p=0,015) in the ipsilateral striatum (Fig. 4B). In the amygdala, NADH-driven FCR was higher in corticosterone conditions (F_*corticosterone*_(1,29) = 5,907, p=0,021, Fig. 4C) without significant differences in uncoupled, succinate-driven FCR (Fig. 4D).

**Fig. 4:**
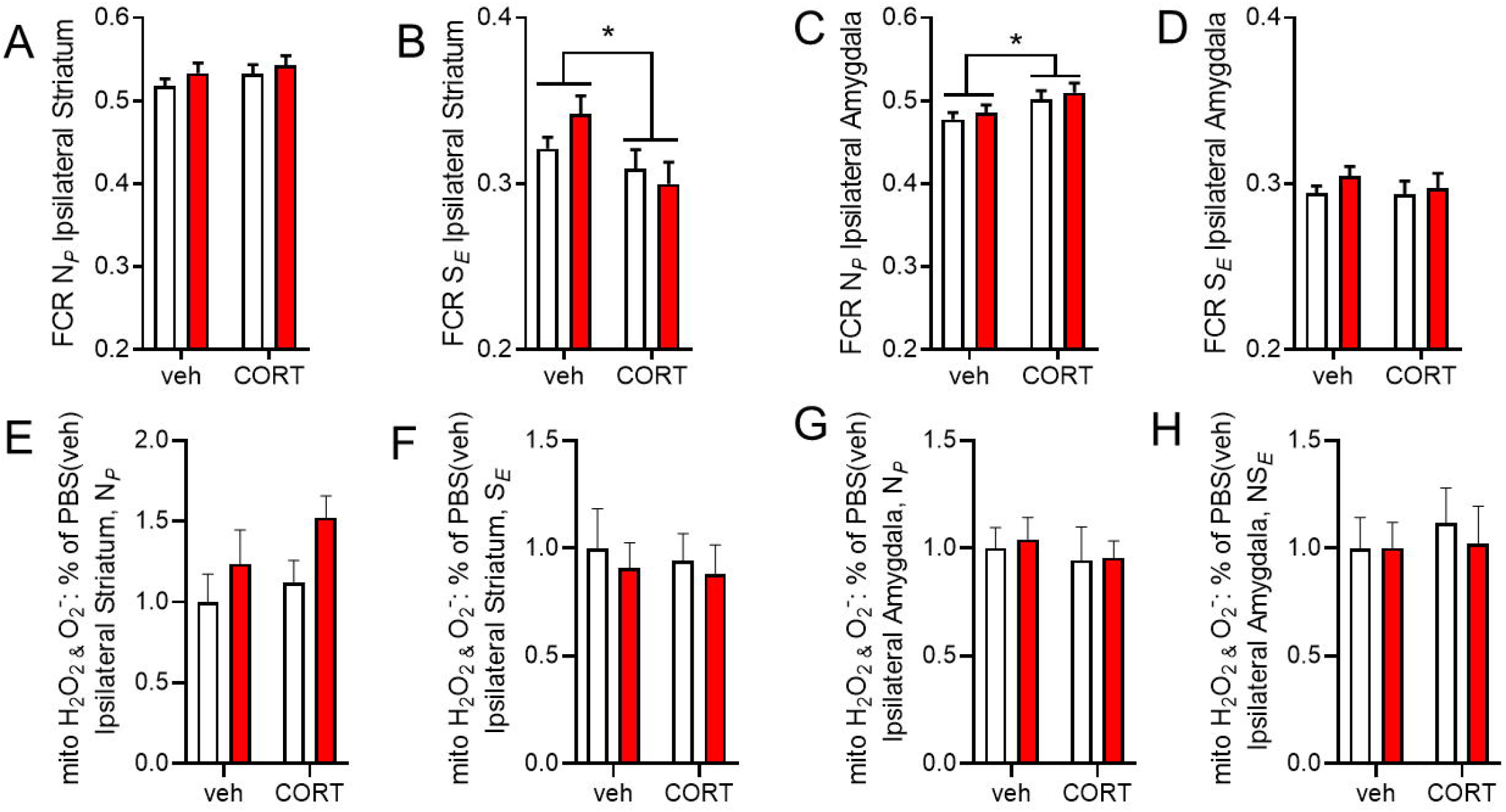
Flux control ratios and mitochondrial ROS. Flux control ratios (FCRs) are presented for striatum (A,B) and amygdala (C,D). Mitochondrial ROS production was similar across groups both in striatum (E,F) and amygdala (G,H). Main effects for CORT are indicated: * p<0.05

We hypothesized that the expected hyperactivity of the amygdala following chronic corticosterone treatment, or α-synuclein pathology might result in increased mitochondrial ROS generation. However, amplex red fluometry revealed no differences in mitochondrial ROS production either in the striatum (Fig. 4E,F) or amygdala (Fig. 4G,H).

## Discussion

Following up on our findings that α-synuclein pathology in mouse amygdala did not cause strong behavioral deficits (Burtscher et al., 2019), we interrogated here, whether α-synuclein pathology, as defined above, was sufficient to cause mitochondrial dysfunction in the amygdala or at the site of injection (striatum). In the applied model, histological staining of pS129 α-synuclein suggests pathology to peak between 1-3 months in the amygdala and decrease thereafter. Conversely, in the striatum the pathology does not decrease, but changes from predominantly neuritic localization at the investigated time-point to successively more perinuclear localization at later time point (Burtscher et al., 2019). As in the present study, we were interested in the interplay of mitochondrial dysfunction and α-synuclein pathology at the time point where pS129 α-synuclein immunoreactivity peaks in the amygdala, we cannot rule out progressive mitochondrial dysfunction at later time points in this model, e.g. in the striatum.

Impaired electron transport chain function due to aggregated forms of α-synuclein has been repeatedly reported in cellular models (Chinta et al., 2010; Reeve et al., 2015) (Tapias et al., 2017; Kim et al., 2018; Wang et al., 2019), especially at late stages of aggregation formation (Mahul-Mellier et al., 2019), while functional assessment of mitochondria in *in-vivo* models of pathology seeding remains lacking. Therefore, we aimed to study mitochondrial respiration directly on fresh brain tissue from regions affected by α-synuclein pathology using *ex-vivo* respirometry, the gold standard to assess mitochondrial function.

We did not observe differences in absolute respiration related to α-synuclein pathology both in the amygdala and the striatum, while chronic corticosterone treatment (associated with clear depression-like phenotypes, but not with significantly changed α-synuclein pathology (Burtscher et al., 2019)) increased mitochondrial respiration in the amygdala. The observation of elevated respiration is in line with previous reports that systemic administration of corticosterone can increase the activity of the basolateral amygdala (Kavushansky and Richter-Levin, 2006). The respirometry protocols applied here furthermore allowed the identification of primarily NADH-linked substrates to drive this increased respiration.

To investigate potential qualitative changes in respirational patterns, we calculated FCRs (Gnaiger, 2009), and detected a reduced succinate-driven FCR in corticosterone conditions in the striatum. Higher succinate-driven FCR after the surgical intervention in the absence of exogenous corticosterone might be an adaptation response of the damaged tissue (Lukyanova and Kirova, 2015; Weilnau et al., 2018). Protein aggregation is believed to induce conditioning-like cellular adaptations (Mao and Crowder, 2010). Liu and colleagues reported less efficient pre-conditioning related adaptations, if animals were treated with exogenous corticosterone (Liu and Zhou, 2012) and the HPA(hypothalamus – pituitary gland – adrenal gland)-axis has been shown to be intimately linked with molecular adaptations to stressors (Rybnikova and Samoilov, 2015), which might explain the absence of succinate-driven FCR upregulation in the corticosterone condition. NADH-driven FCR in the oxidative phosphorylation state was higher in the corticosterone conditions in the amygdala, confirming the focal role of complex I in the observed effect of enhanced respiration after chronic corticosterone.

Misfolded α-synuclein has previously been associated with oxidative stress (Hsu et al., 2000). Also chronic corticosterone treatment has been demonstrated to result in a dysregulated oxidative balance (Liu and Zhou, 2012). Therefore, we asked, whether α-synuclein pathology or chronic corticosterone resulted in increased mitochondrial ROS production in our model, which was not the case. In summary, these results suggest that α-synuclein pathology as defined here (immunoreactivity to pS129 and C-terminal truncated, insoluble forms of α-synuclein) and 5-6 weeks post injection is not sufficient to induce clear mitochondrial dysfunction in mouse amygdala and striatum. Additional challenge of amygdala circuits by corticosterone did not result in mitochondrial dysfunction, although it might have blocked mitochondrial adaptations in response to α-synuclein pathology formation. A summary of the reported findings is given in Fig. 5.

**Fig. 5:**
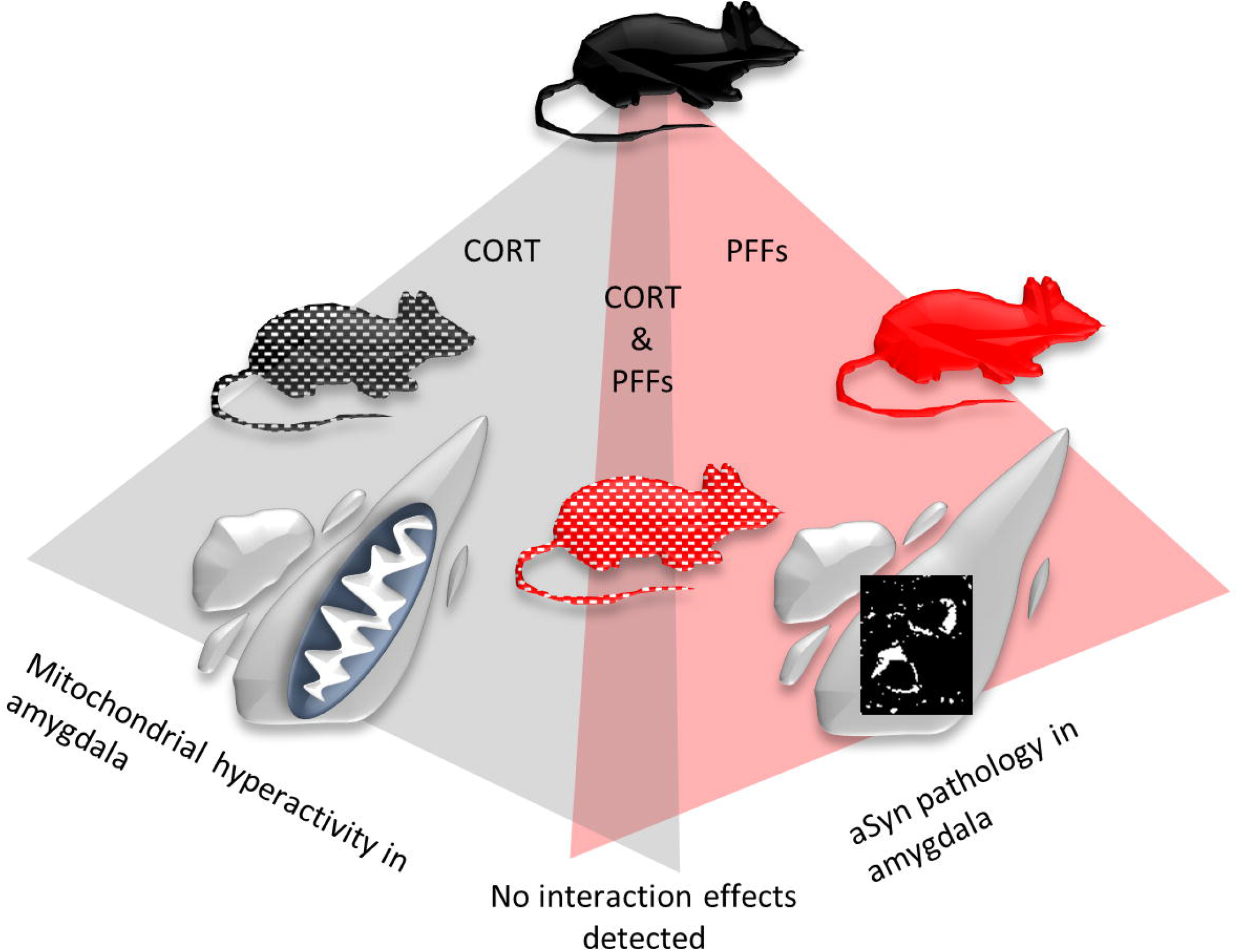
Working model. While chronic corticosterone induced respirational hyperactivity in the amygdala and PFF inocculation resulted in pronounced α-synuclein (α-synuclein) pathology, no deficits of mitochondrial respiration were observed due to α-synuclein pathology.

Importantly, as we did not observe pS129 α-synuclein in striatum or amygdala 5-6 weeks after PFF injection, even in the insoluble fractions, we assume that at this time-point pS129-positive, insoluble aggregates are too rare to be detected by western blotting. Conversely, by immunostaining pS129 α-synuclein was readily detected. This could mean that pS129-positive α-synuclein pathology precedes the formation of insoluble aggregates, which is line with our recently presented study on the successive maturation of pS129 α-synuclein-positive aggregates into Lewy body like structures (Mahul-Mellier et al., 2020).

A limitation of the here performed approach is the possibility that mitochondrial dysfunction was not detected, because of a too low number of affected neurons. While we cannot exclude this possibility, which warrants further investigation, we demonstrate that overall brain region respirational patters were maintained also in the presence of dense pS129 α-synuclein pathology. On the other hand, chronic supply of exogenous corticosterone increased mitochondrial respiration in the amygdala independently of pS129 α-synuclein pathology.

## Conclusions

Both α-synuclein pathology and mitochondrial dysfunction have been implicated in the pathogenesis of PD and other synucleinopathies. Despite recent advances in the elucidation of the molecular underpinnings of both processes, it remains unclear, how α-synuclein pathology influences mitochondrial function *in vivo* and vice versa and if one or both of these hallmarks are sufficient to cause neurodegeneration and the manifestation of clinical symptoms. Our results suggest that the prominent α-synuclein pathology observed at 5-6 weeks post-inoculation with PFFs and as defined above is not sufficient to induce mitochondrial dysfunction. Although significant cell loss in the substantia nigra in this model is usually only detected 6 months after intrastriatal PFF injection (Luk et al., 2012), we expected the pronounced α-synuclein pathology early after seeding to have effects on mitochondrial function. A recent report demonstrated a non-significant trend towards reduced neuron numbers in the basolateral amygdala also only 6 months, and only after bilateral injection of high PFF concentrations (Stoyka et al., 2019).

Our findings suggest that mitochondrial dysfunction may occur later, possibly during post-α-synuclein aggregation events linked to the transition from pS129 immunoreactive filamentous aggregates to LBs, which also involve the recruitment of mitochondria and other membranous organelles into LBs. Indeed, recent studies in primary neuron models have shown that recruitment of mitochondria and mitochondrial dysfunction occur primarily at later stages of LB formation and maturation (Mahul-Mellier et al., 2019). Together, these findings highlight the critical importance of revisiting the interplay between α-synuclein and mitochondria at the various stages on the pathway to LB formation and their roles in LB formation and maturation and neurodegeneration in this model and other animal models of PD pathology and synucleinopathies.

Our findings underscore caution to be taken regarding the translational validity of this model, given the weak symptomatology, in particular of non-motor symptoms at the peak of α-synuclein pathology in this model (1-3 months after injection of PFFs)(Burtscher et al., 2019), and difficult to reproduce motor-symptoms, even at late stages of pathology, and even by the leading groups in the field, For example, recently Henderson et al reported no deficits in rotarod-assessed motor coordination (Henderson et al., 2019) even up to 9 months after PFF-injection, despite this being one of the most replicated feature in this model (Luk et al., 2012; Kim et al., 2019). We also warn about potential publication bias, with negative reports on this model becoming insufficiently public and encourage other scientist to share and discuss their negative findings.

## Acknowledgements

We are grateful to Olivia Zanoletti for excellent technical support and to Somanath Jagannath and Salvatore Novello for constructive feedback. This work was funded by EPFL and UCB S.A.

## Author’s roles

JB, JCC, CS and HAL contributed to the research and statistical design. JB and JCC performed the experiments and statistical analysis. JB wrote the first draft of the manuscript. JB and HAL wrote the manuscript. JCC and CS reviewed the manuscript.

